# Too packed to change: site-specific substitution rates and side-chain packing in protein evolution

**DOI:** 10.1101/013359

**Authors:** María Laura Marcos, Julian Echave

## Abstract

In protein evolution, due to functional and biophysical constraints, the rates of amino acid substitution differ from site to site. Among the best predictors of site-specific rates is packing density. The packing density measure that best correlates with rates is the weighted contact number (WCN), the sum of inverse square distances between the site’s *C*_*α*_ and the other *C*_*α*_. According to a mechanistic stress model proposed recently, rates are determined by packing because mutating packed sites stresses and destabilizes the protein’s active conformation. While WCN is a measure of *C*_*α*_ packing, mutations replace side chains, which prompted us to consider whether a site’s evolutionary divergence is constrained by main-chain packing or side-chain packing. To address this issue, we extended the stress theory to model side chains explicitly. The theory predicts that rates should depend solely on side-chain packing. We tested these predictions on a data set of structurally and functionally diverse monomeric enzymes. We found that, on average, side-chain contact density (WCN_*ρ*_) explains 39.1% of among-sites rate variation, larger than main-chain contact density (WCN_*α*_) which explains 32.1%. More importantly, the independent contribution of WCN_*α*_ is only 0.7%. Thus, as predicted by the stress theory, site-specific evolutionary rates are determined by side-chain packing.

## INTRODUCTION

Why do some protein sites evolve more slowly than others? Protein evolution is driven by random mutations and shaped by natural selection (Liberles et al., 2012; Sikosek and Chan, 2014). Mutations are selected depending on their impact on functional properties, such as the chemical nature of catalytic residues, active site conformation, and the protein’s ability to fold rapidly and stably. Since changes of these properties depend on the mutated site, amino acid substitution rates vary from site to site.

We can reformulate the question opening the previous paragraph: What *specific properties* account for site-dependent rates of evolution? Substitution rates have been found to correlate with several properties. Amongst the best predictors are solvent accessibility (Bustamante et al., 2000; Conant and Stadler, 2009; Franzosa and Xia, 2009; Ramsey et al., 2011; Shahmoradi et al., 2014) and stability changes (Echave et al., 2014). For a large data set of enzymes, it was found that the main structural determinant is the *weighted contact number* WCN (Shih and Hwang, 2012; Yeh et al., 2014a,b). A site’s WCN is the sum of inverse square distances from its *C*_*α*_ to the *C*_*α*_s of other sites, therefore, it is a measure of *C*_*α*_ packing density.

The relationship between WCN and substitution rates can be understood in terms of a mechanistic stress model of protein evolution (Huang et al., 2014). Given an ancestral wild-type protein, the model assumes that its native conformation is the active conformation. Mutating a site perturbs (stresses) its interactions with other sites, destabilizing the active conformation. Such a destabilization determines the probability of the mutation being accepted or rejected, and therefore the rate of amino acid substitutions. Using the parameter-free Anisotropic Network Model (Yang et al., 2009), the expected destabilization was found to be proportional to WCN, and site-specific substitution rates were predicted to decrease linearly with increasing WCN, in agreement with observations.

So far, substitution rate vs. WCN studies were based on main chain (*C*_*α*_) packing (Shih and Hwang, 2012; Yeh et al., 2014a; Huang et al., 2014). However, mutations replace side chains. Consider a protein residue, e.g. Thr93 of Human Carbonic Anhidrase II (pdb code 1CA2) (Fig. 1). The environment of the main chain (panel A) differs from that of the side chain (panel B). When Thr93 is mutated, what environment would determine whether the mutation is accepted or rejected? More specifically: Do site-specific substitution rates depend on main-chain packing or on side-chain packing? To address this issue, we extended the stress model to consider main and side chains explicitly, we derived substitution rates as a function of packing, and tested the theory on a data set of monomeric enzymes.

**Figure 1.**
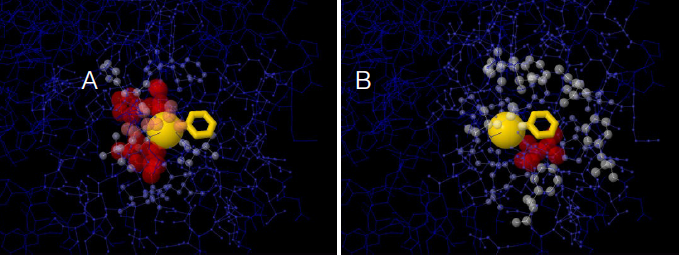
The two environments of a protein residue. Images of the environments of Thr93 of Human Carbonic Anhidrase II (pdb code 1CA2). (A) Environment of the main chain *C*_*α*_: the size and colors of protein atoms increase with the inverse square distance to Thr93 *C*_*α*_ (gold ball). (B) Environment of the side chain: size and colors of atoms increase with the inverse square distance to the geometric center of Thr93 side chain (gold wireframe).

## METHODS

### The stress model

The stress model provides a mechanism for the observed correlation between rates and packing density (Huang et al., 2014). The model is based on the idea that a mutant is viable to the extent that it spends time in the active conformation. When a site is mutated, the interactions with its neighbors are perturbed (stressed), which destabilizes the active conformation by an amount *δV*\*, the *local mutational stress*. Mutational stress is related to site-specific evolutionary rates:

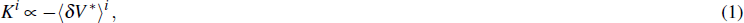

i.e. the substitution rate of site *i*, *K*^*i*^, decreases linearly with the *mean local mutational stress*, 〈*δV*\*〉^*i*^ (*δV*\* averaged over mutations at *i*). (1) is the main equation of the stress theory.

To calculate the mutational stress, we need an energy function. Huang et al. (2014) used the *parameter-free Anisotropic Network Model* (pfANM) of (Yang et al., 2009), which models the protein using an elastic network where each residue is represented by a node placed at its *C*_*α*_. Pairs of nodes are connected by springs with force constants 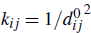, where 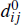 is the distance between 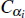 and 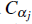 in the active conformation. Following (Echave, 2008; Echave and Fernández, 2010), mutations are modeled as random perturbations of the lengths of the springs connected to the mutated site, which leads to:

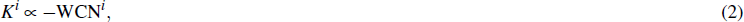

where

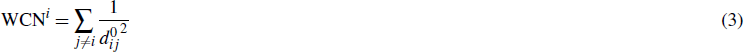

is the weighted contact number introduced by Lin et al. (2008) and found to be among the best structural predictors of site-dependent evolutionary rates (Yeh et al., 2014a,b). Thus, according to the stress theory combined with the *C*_*α*_-based pfANM, substitution rates should decrease linearly with WCN.

Since point mutations replace *side chains*, including them explicitly might improve the predictions of the stress theory. To explore this possibility, we model the protein as an elastic network where each residue is represented by two nodes, one for the main chain, *α*, placed at the residue’s *C*_*α*_, and another for the side chain, *ρ*, placed at the side-chain geometric center (only *α* nodes for Glycines). Mutations affect only the side chain of the mutated site. We model them adding random perturbations to the lengths of the springs connected to the mutated site. Assuming, as before, that the force constant of the spring connecting nodes *n*_*i*_ and *n*_*j*_ (*n* is *α* or *ρ*) is 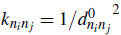, it follows that:

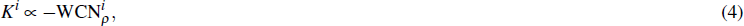

where

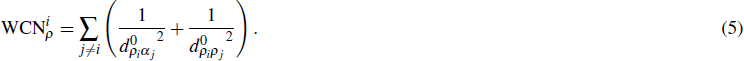

WCN_*ρ*_, defined here, is the weighted contact number of the side chain. Thus, when using the pfANM based on main chain nodes *α* and side-chain nodes *ρ*, the stress model predicts that site-specific rates will depend only on the contact density of the side chain WCN_*ρ*_. To check this prediction, we also consider the main chain weighted contact number:

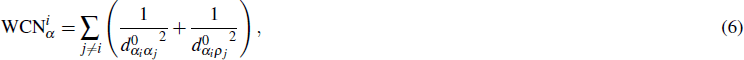

According to the stress model, main-chain packing should not contribute independently to substitution rates.

In this section, we briefly presented the main results of the stress theory. A more detailed derivation can be found in the Appendix and in (Huang et al., 2014).

### Dataset and comparison of empirical and predicted rates

To test our theory, we used the data set of (Echave et al., 2014). The set consists of 209 monomeric enzymes of known structure covering diverse structural and functional classes. Each structure is accompanied by up to 300 homologous sequences.

We used the empirical site-specific rates of evolution of (Echave et al., 2014). They were calculated as follows. First, the homologous sequences for each structure were aligned using MAFFT (Multiple Alignment using Fast Fourier Transform) (Katoh et al., 2005; Katoh and Standley, 2013). Second, using the resulting alignments as input, Maximum Likelihood phylogenetic trees were inferred with RAxML (Randomized Axelerated Maximum Likelyhood), using the LG substitution matrix (named after Le and Gacuel) and the CAT model of rate heterogeneity (Stamatakis, 2014). Third, the alignment and phylogenetic tree for each structure was used as input of Rate4Site to obtain the site-specific rates of substitution using the empirical Bayesian method and the amino-acid Jukes-Cantor mutational model (aaJC) (Mayrose et al., 2004). Finally, site-specific *relative* rates were obtained by dividing site-specific rates by their average over all sites of the protein. We denote the empirical rates by *K*_R4S_.

For each protein, we calculated three packing density measures and predicted rates using linear fits. For brevity, we will use the shorthand *y* ∼ *x* for one-variable linear fits and *y* ∼ *x*_1_ + *x*_2_ for two-variable fits. Using the protein’s pdb structure, we calculated WCN, WCN_*α*_, and WCN_*ρ*_, using (3), (6), and (5), respectively. Then, we calculated predicted rates by fitting *K* ∼ WCN, *K* ∼ WCN_*α*_, and *K* ∼ WCN_*ρ*_ to the set of empirical rates. We also considered the two-variable fit *K* ∼ WCN_*α*_ + WCN_*ρ*_. The goodness of fit of each model was assessed using *R*^2^, the square correlation coefficients between predicted and empirical rates.

For statistical analysis we used R (R Core Team, 2014). For linear fits we used the built-in function `lm()`. Correlation coefficients were calculated using `cor()`. Binomial tests were performed `binom.test()`.

## RESULTS AND DISCUSSION

According to the stress model, site-specific substitution rates depend only on side-chain packing. Main chain packing should not be directly related to substitution rates. To test this theory, we compared rate predictions based on main-chain packing and side-chain packing for a data set of 209 diverse monomeric enzymes.

Consider, for example, Human Carbonic Anhidrase II (pdb code 1CA2). Empirical rates *K*_R4S_ were obtained from the multiple sequence alignment as described in Methods. Using the pdb structure, we calculated the packing measures WCN, WCN_*α*_, and WCN_*ρ*_, using (3), (6), and (5), respectively. We used these packing measures to predict rates using linear fits to empirical rates, as described in Methods. As we mentioned in the Introduction, main chain environments and side-chain environments are different (Fig. 1). Accordingly, WCN_*α*_ and WCN_*ρ*_ result in different predicted rates (Fig. 2). The two site-dependent profiles of predicted rates are similar to the empirical *K*_R4S_ profile. WCN_*ρ*_-based predictions look better (Fig. 2) and are better (Fig. 3): the *R*^2^ values are 0.41 for WCN_*α*_ and 0.56 for WCN_*ρ*_. *R*^2^ increases only by 0.02 for the two-variable fit *K* ∼ WCN_*α*_ + WCN_*ρ*_ (*R*^2^ = 0.58). Since WCN (Eq. (3)) was, so far, the best structural predictor of site-specific rates for enzymes (Yeh et al., 2014a,b), we also calculated *R*^2^(*K*_R4S_, WCN): it is 0.40. To summarize, for 1CA2, WCN_*ρ*_ > WCN_*α*_ ≳ WCN; the best predictor of site-specific rates is WCN_*ρ*_. Moreover, WCN_*α*_ has only a small independent effect on substitution rates.

**Figure 2.**
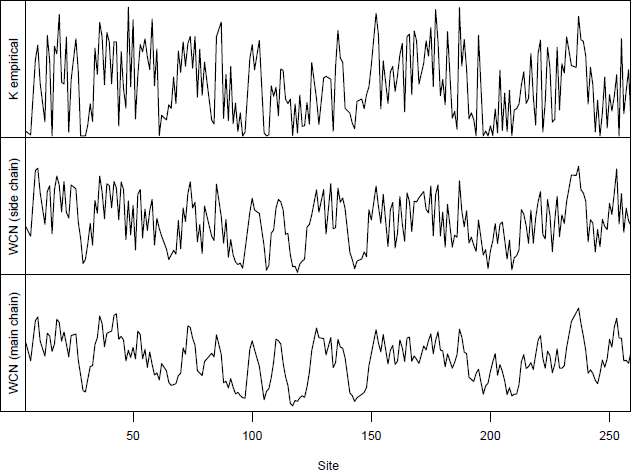
Profiles of site-specific evolutionary rates for 1CA2. (Top) empirical rates *K*_R4S_ inferred by Rate4Site. (Middle) Rates predicted from the side-chain contact density WCN_*ρ*_. (Bottom) Rates predicted from the main-chain contact density WCN_*α*_. The profile of WCN_*ρ*_-predicted rates looks more similar to the *K*_R4S_ profile

**Figure 3.**
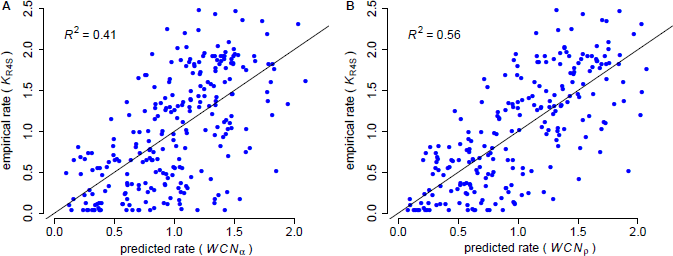
Empirical vs. predicted rates for 1CA2. (A) Empirical rates inferred using Rate4Site vs. rates predicted from the main-chain contact densities WCN_*α*_. (B) Empirical rates vs. rates predicted from side-chain contact densities WCN_*ρ*_. The “x=y” line corresponding to a perfect fit is shown. WCN_*α*_ explains *R*^2^ = 41% of the variation of site-specific empirical rates, WCN_*ρ*_ explains 56%.

We repeated the previous assessment for each protein of the data set. For each of the 209 enzymes, we calculated the densities WCN, WCN_*α*_, and WCN_*ρ*_ and calculated predicted rates from linear fits to empirical rates *K*_R4S_. *R*^2^ values averaged over all proteins are 0.316, 0.321, and 0.391 for WCN, WCN_*α*_, and WCN_*ρ*_, respectively (Fig. 4). Thus, as for 1CA2, the predictive power of single-variable fits follows WCN_*ρ*_ > WCN_*α*_ ≳ WCN. When going from *K* ∼ WCN_*ρ*_ to *K* ∼ WCN_*α*_ + WCN_*ρ*_, *R*^2^ increases from 0.391 to 0.398, only a 0.7% increase in explained variance (Fig. 4). Therefore, on average, WCN_*ρ*_ is the best predictor of site-specific substitution rates.

**Figure 4.**
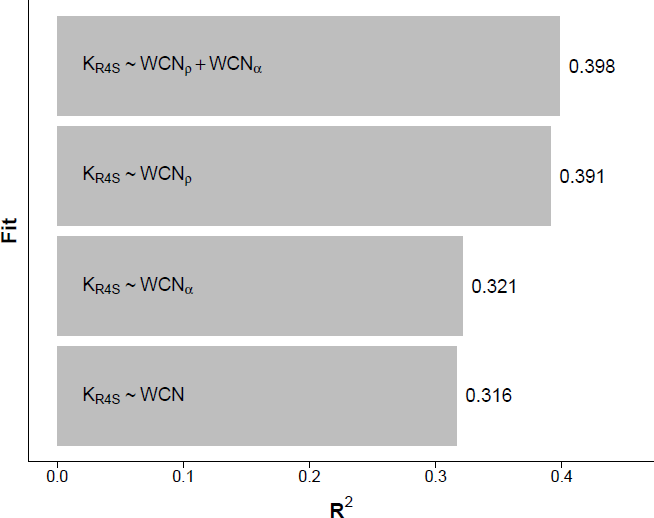
Side chain packing is the sole determinant of site-specific substitution rates. Linear regression of empirical rates (*K*_R4S_) using three one-variable fits (WCN, WCN_*α*_, and WCN_*ρ*_) and a two variable fit *K*_R4S_ ∼ WCN_*α*_ + WCN_*ρ*_. WCN and WCN_*α*_ are measures of the contact density of main chain *C*_*α*_s. WCN_*ρ*_ is the contact density of side chains, modeled by their geometric centers. The models are fit for each protein, and *R*^2^ is the average *R*^2^ over the 209 proteins of the data set. WCN_*ρ*_ is the best predictor. The independent contribution of WCN_*α*_is very small (0.7%).

Beyond average *R*^2^, we performed a protein-by-protein comparison (Fig. 5). We found that WCN_*ρ*_ is a better predictor than WCN_*α*_ for 204 of the 209 proteins studied (*p* ≪ 10^−3^, binomial test). Similarly, WCN_*ρ*_ outperforms WCN for 206/209 proteins (*p* ≪ 10^−3^, binomial test). Thus, WCN_*ρ*_ is the best rate predictor for almost all proteins of the data set.

**Figure 5.**
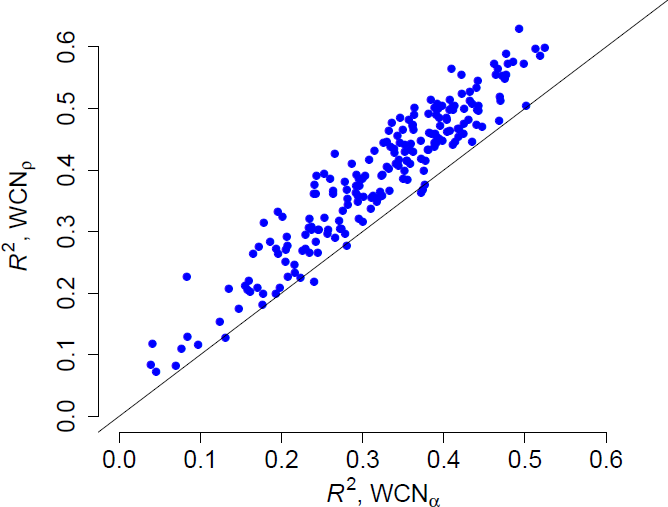
Side chain packing is the best predictor of substitution rates for most proteins. *R*^2^ is the square correlation between empirical rates (*K*_R4S_) and either side-chain contact density WCN_*ρ*_ (*y* axis) or main-chain contact density WCN_*α*_ (*x* axis). Each point corresponds to one protein. Empirical rates correlate better with WCN_*ρ*_ for 204 out of 209 proteins.

To summarize, side-chain contact density (WCN_*ρ*_) is the best predictor of site-specific substitution rates, accounting, on average, for 39.1% of the rate variation among sites. In contrast, the independent contribution of main-chain contact density (WCN_*α*_) is negligible (0.7%). These results are consistent with the predictions of the stress model, extended to include explicitly main chain and side chains. According to this theory, mutations replace side chains thus changing the parameters of interaction between the mutated side chain and the rest of the protein. WCN_*ρ*_ is proportional to the destabilization of the protein’s active conformation, which is why it correlates with rates: mutations are accepted or rejected according to the degree of destabilization of the active conformation.

From a practical point of view, regardless of the validity of the stress theory, WCN_*ρ*_ outperforms WCN, that was, so far, the best structural predictor of site-specific substitution rates (Yeh et al., 2014a,b). Therefore, at least for the data set of monomeric enzymes used, WCN_*ρ*_ is the new best predictor of site-specific substitution rates. WCN_*ρ*_ could be used to improve structure-based empirical models of protein evolution and phylogenetic inference (see e.g. (Kleinman et al., 2010)).

## APPENDIX

### The stress model

The stress model is based on the idea that a mutant is viable to the extent that it spends time in the active conformation. The fixation probability is modeled as

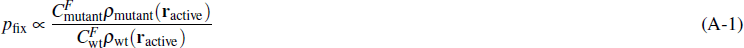

where *C*^*F*^ is the concentration of folded protein and *ρ* (**r**_active_) is the probability of it adopting the active conformation. Assuming that *C*_mutant_/*C*_wt_ is equal to the ratio of partition functions, from basic statistical physics it follows that:

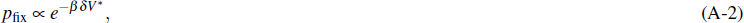

where *β* can be thought of representing selection pressure rather than temperature and

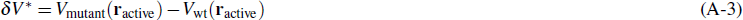

is the energy difference between mutant and wild-type in the active conformation. Finally, assuming that *βδV*\* ≪ 1 (weak selection), from (A-2) we find:

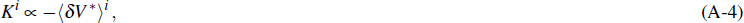

i.e. the rate of substitution of site *i*, *K*^*i*^, is proportional to (minus) the destabilization energy averaged over mutations at *i*, 〈*δV*\*〉^*i*^. This is the basic equation of the stress theory.

### One-bead-per-site elastic network

To derive substitution rates, we need an energy function. Let us model the protein as an elastic network of nodes placed at *C*_*α*_s connected by elastic springs. The energy of a conformation **r** is given by:

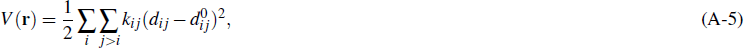

where *d*_*ij*_ = ∥**r**_*j*_ − **r**_*i*_∥ is the distance between 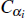 and 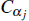, *k*_*ij*_ is the force constant of spring *i* − *j* and 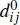 its equilibrium length.

The wild-type protein is modeled by using springs that are relaxed at the native conformation: 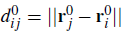. To model a mutation at site *i*, we add random perturbations to the spring lengths connecting *i* other sites: 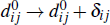. Using (A-5) and (A-3), we find:

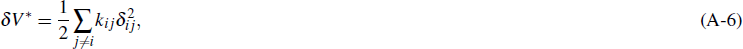

where we have assumed that the active conformation is the native conformation of the wild type, 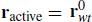, which is a reasonable assumption for purifying selection. Assuming that *δ*_*ij*_ for the different contacts are drawn independently from the same distribution, averaging (A-6) over mutations at site *i* we find

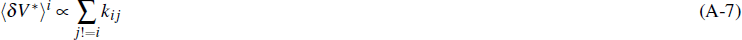

Thus, the mutational destabilization averaged over mutations at a given site (the *mean local mutational stress*) is proportional to the sum of the force constants of the springs connected to the mutated site.

To obtain the site-specific rates, we use the parameter-free Anisotropic Network Model (pfANM):

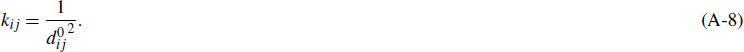

Replacing (A-8) into (A-7), and the result into (A-4), we obtain:

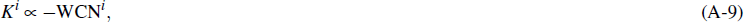

where

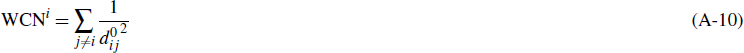

is the weighted contact number. Thus, when using the pfANM based on *C*_*α*_, the stress model predicts that site-specific rates are proportional to (minus) WCN.

### Two beads-per-site elastic network

The elastic network of the previous section uses one node per site, thus modeling side chains only implicitly. Since mutations replace side chains, including them explicitly might improve the predictions of the stress theory.

Let us represent each site using two nodes: one for the main chain, *α*, placed at the residue’s *C*_*α*_ as before, and another for the side chain, *ρ*, placed at the side-chain geometric center (Gly’s are represented using only one node at *C*_*α*_). The elastic energy is:

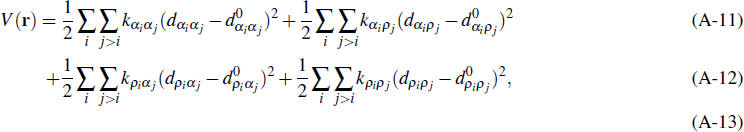

where 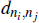 is the distance between nodes *n*_*i*_ and *n*_*j*_ (*n* is *α* or *ρ*), 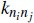 is the force constant of the spring connecting these nodes, and 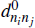 the equilibrium spring length.

A mutation at site *i* will replace *ρ*_*i*_, affecting only the parameters of the energy function related to this node. Modeling a mutation at *i* by adding random perturbations to the springs of 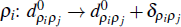 and 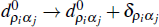, we find:

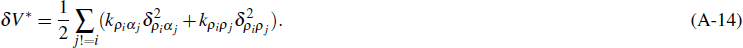

where *δV*\* is defined in (A-3). Assuming perturbations are drawn independently from the same distribution, averaging (A-14) over mutations at *i* we find:

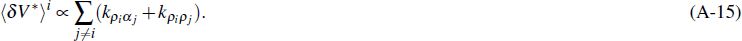

Finally, assuming that 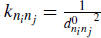, from (A-4) and (A-15) we obtain:

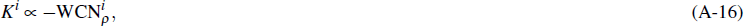

where

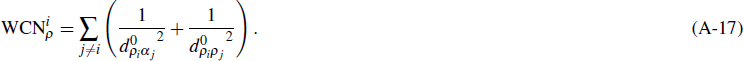

WCN_*ρ*_, defined here, is the side-chain weighted contact number. Thus, when using the pfANM based on main chain nodes *α* and side-chain nodes *ρ*, the stress model predicts that site-specific rates will depend on the contact density of the side chain WCN_*ρ*_.

For the sake of the present study, we also consider whether main-chain packing has any independent effect on rates. For this purpose, we calculate

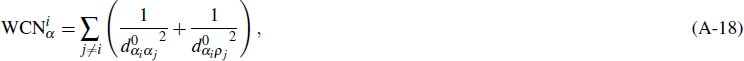

which is the main chain weighted contact number for the two-beads-per-site network model.

